# ARG-PASS v2.0 expands global resistome surveillance to reveal previously unrecognized antibiotic resistance genes reaching the human gut

**DOI:** 10.64898/2026.10.21.760772

**Authors:** Lachlan Bartrop, Frédéric Grenier, Joseph Bielawski, Sébastien Rodrigue, Louis-Patrick Haraoui

**Affiliations:** Department of Microbiology and Infectious Diseases, Université de Sherbrooke, Québec, Canada; Department of Biology, Université de Sherbrooke, Québec, Canada; Department of Biology, Dalhousie University; Centre de recherche Charles-Le Moyne, CISSS Montérégie-Centre, Greenfield Park, Québec, Canada.; Humans & the Microbiome Program, Canadian Institute for Advanced Research, Toronto, Ontario, Canada

## Abstract

Antimicrobial resistance (AMR) is a significant global public health threat, and antibiotic resistance genes (ARGs) that bacteria depend on to withstand treatment are its molecular basis. How ARGs move from environmental reservoirs into the human microbiome, and eventually into human pathogens, remains one of the most important unresolved questions in AMR research. The environmental reservoir remains poorly described and the routes connecting it to human microbiomes are largely inferred rather than observed. Describing that reservoir requires computational methods able to recognise ARGs that share little similarity with known enzymes, yet retain accuracy after experimental validation to enable scalable applications. Dietary exposure through unprocessed ferments offers a tractable setting in which such a route can be observed directly. Here we introduce ARG-PASS v2.0 and first apply it to a lake metagenome from Kolkata, India, where 10 of 11 tested candidates across diverse ARG classes were functionally confirmed after expression in *Escherichia coli*. Given their clinical relevance, we then predicted 918 class B1 metallo-β-lactamases (MBLs) from MGnify and the PLSDB, with most diverging from characterized enzymes, but as widely distributed. Often tracking their hosts’ distribution, we find that most class B1 MBL diversity lies outside human-associated microbiomes. Cross-referencing against the Unified Human Gastrointestinal Protein catalogue identified a carbapenemase, an ANT, and class A β-lactamases from *Bacillus sonorensis* and *Bacillus licheniformis*, detected in South Korean fermented soybean paste (*doenjang*) as well as gut isolates of populations consuming these foods. Together, these results point to a largely chromosomal reservoir of ARG diversity that reaches the human gut through both resident and generalist hosts. Once in the human microbiome, treatment may be compromised directly if the host is an opportunistic pathogen, or after transfer to a clinical pathogen.

## Introduction

Antibiotics form the foundation of modern medicine, yet their utility is threatened by antimicrobial resistance (AMR), projected to become a leading cause of death globally by mid-century (O’Neill, 2016). Antibiotic resistance genes (ARGs) cause AMR, but how they transition from environmental reservoirs to the human microbiome, and ultimately to clinical pathogens, remains one of the most important unresolved questions in AMR research. Answering it requires knowing which ARGs exist, where they reside, in which hosts, and by what route they might reach clinical settings. Surveillance has largely focused on genes already associated with mobile genetic elements or resembling those characterised clinically, so the environmental reservoir remains poorly described, and the routes by which it connects to human microbiomes are largely inferred rather than observed (Larsson & Flach, 2022). Dietary exposure is one such route; unprocessed ferments deliver substantial quantities of viable microorganisms directly to the gastrointestinal tract (Derrien & Vlieg, 2015; Wolfe, 2023), and antibiotic-resistant *Klebsiella* have recently been characterized across Chinese fermented soybean products with evidence of possible lateral gene transfer (LGT) to gut microbiomes (Xiang et al., 2025). However, whether functionally verified, previously uncharacterized (novel) ARGs reach the gut by this route remains undemonstrated.

Practical resistome surveillance depends on identifying novel ARGs before they become a clinical problem: informing governments, universities, and the pharmaceutical industry of incoming threats, and revealing how ARGs are mobilized and spread by mobile genetic elements (MGEs). ARG detection methods fall broadly into two groups which trade precision for sensitivity. Alignment-based approaches achieve high precision but are restricted to sequences with high identity to known ARGs (Alcock et al., 2023). For example, *sul4* was first identified in polluted river sediment in India but had already disseminated across Asia and Europe (Razavi et al., 2017) and has since been detected in clinical *Salmonella enterica* (Peng et al., 2023). At less than 34% amino acid sequence identity to known sul genes, it would have been missed by alignment-based approaches. Profile Hidden Markov models (Berglund et al., 2019), deep learning (Arango-Argoty et al., 2018; Wu et al., 2023) and protein structure prediction (Ruppé et al., 2019), enhance sensitivity, but are less able to distinguish true novel ARGs from non-ARG homologues (Inda-Díaz et al., 2023). Because phenotypic validation is required before a candidate enters curated databases such as the Comprehensive Antibiotic Resistance Database (CARD), prediction accuracy determines how quickly a determinant becomes available to clinical diagnostics. Moreover, ARGs do not respect enivronmental–clinical boundaries, so understanding how they reach human pathogens requires tracking them across both. Characterising where novel ARGs reside and how they move therefore requires tools that combine sensitivity to divergent sequences with the precision to scale across global metagenomic datasets.

Here we introduce ARG-PASS v2.0 builds on the original method (Bartrop et al., 2026) to detect ARGs with low sequence identity to known ARGs with high precision. When applied to a lake metagenome from Kolkata, India, 10 of 11 predicted candidates across diverse ARG classes were functionally confirmed after heterologous expression in *E. coli*. Given their clinical relevance, we then predicted 918 class B1 metallo-β-lactamases from MGnify and the PLSDB, with most diverging from characterized enzymes (mean CARD identity of 53%), yet no less geographically widespread. We highlight ∼100 candidates which occur in three or more countries and likely track their hosts’ distribution, with some recovered only from human-associated biomes, however most span a wider ecological range. Cross-referencing against the Unified Human Gastrointestinal Protein catalogue identified a class B1 MBL, an ANT and class A β-lactamases from *Bacillus sonorensis* and *Bacillus licheniformis*, detected in South Korean fermented soybean paste (*doenjang*) as well as gut isolates of populations consuming these foods. All doenjang ARGs confer resistance above clinical breakpoints. Together, these results point to a largely chromosomal reservoir of undescribed ARG diversity that reaches the human gut through both resident and generalist hosts, where treatment may be compromised directly or through transfer to clinical pathogens. ARG-PASS v2.0 is freely available to use from GitHub (https://github.com/btroppo/ARG-PASS/)

## Results

### ARG-PASS v2.0 outperforms existing tools for functional ARG prediction

ARG-PASS v2.0 generates a two-dimensional distribution of pairwise sequence identities (seqIDs) vs Template Modelling (TM)-scores of functionally important (conserved) regions of antibiotic resistance protein (ARP) structures and trains a one-class support vector machine (SVM) to learn the decision boundary (DB) surrounding the distribution. A candidate protein structure is aligned to the conserved regions of each ARP structure and its derived seqIDs and TM-scores are calculated. If any points lie within the DB, the candidate is predicted functional; if all points lie outside the DB, it is flagged as an outlier and predicted non-functional (Figure 1a, c). ARG-PASS v2.0 is an advance from the original method (Bartrop et al., 2026), through more precise identification of functionally important residues within each ARG class, achieved using CONSTRUCT (Chivot et al., 2025). For applications requiring greater sensitivity, such as detecting ARG-like folds associated with elevated minimum inhibitory concentrations (MICs) but remain below clinically established breakpoints (i.e., pre-resistance genes), we retain local Distance Difference Test (lDDT) (Bartrop et al., 2026) scoring as an alternative mode, which we refer to as ARG-PASS v2.0 (lDDT). ARG-PASS v2.0 refers to the default configuration throughout unless the lDDT mode is specified.

**Fig 1.**
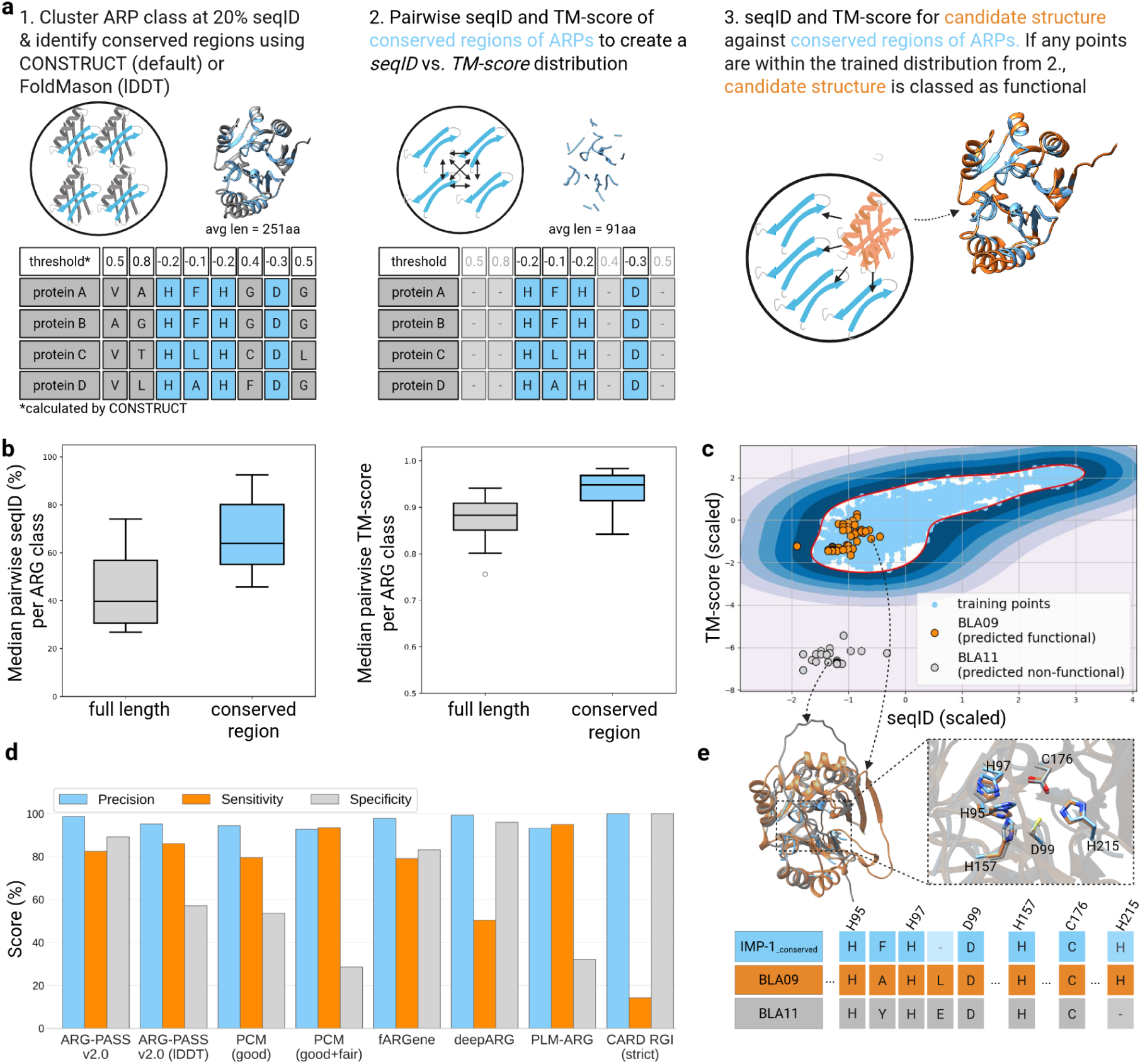
Overview of ARG-PASS v2.0. Step 1. Antibiotic resistance protein (ARP) structures for an ARG class (here as class B1 MBLs, with IMP-1 as an example) are clustered at 20% sequence identity (seqID) using Foldseek. Conserved regions are then extracted from each cluster by applying a threshold to per-residue scores calculated either with CONSTRUCT or as a local distance difference test (lDDT); these two routes define the default and lDDT configurations of ARG-PASS v2.0 respectively. Step 2. The pairwise (all-vs-all) seqIDs vs TM-scores are calculated for conserved regions of ARP structures within clusters using Foldseek TM-align, to create the training distribution. Step 3. The seqIDs and TM-scores comparing a candidate protein structure and conserved regions of known ARP structures are calculated using Foldseek TM-align. A one-class SVM is trained on the *x*, *y* distribution from 2, where the decision boundary (DB, in red) surrounds all training points (c). The seqIDs vs TM-scores of the aligned region of a candidate structure against each conserved ARP structure are plotted (data points in orange). If any data points lie on or within the DB, the candidate structure is classified as functional (c, BLA09 functionally verified in this study is classified as functional), otherwise the candidate is classified as non-functional (c, BLA11, lacked activity and correctly predicted non-functional by ARG-PASS v2.0). b, Maximum seqID and TM-score between ARP structures vs conserved regions of ARP structures (Mann-Whitney test between each condition: p < 0.001). d, Benchmarking results of ARG-PASS v2.0 against existing ARG prediction methods. e, The conserved region of IMP-1 aligned to the functional BLA09 and non-functional BLA11. BLA11 lacks Histidine residue H263 (H215 in the alignment) in the Zn2+ binding site of class B1 MBLs which likely contributed to no phenotype observed. For further details see Methods. Figure created with Biorender.com.

We benchmarked ARG-PASS v2.0 against existing computational methods designed for ARG prediction (PCM, fARGene, deepARG, PLM-ARG) and a reference tool based on sequence similarity (RGI strict), using a manually curated validation dataset of positive (*n* = 280) and negative (*n* = 27) experimentally verified resistance determinants across a large CARD identity range (Fig 1d, Table S2). ARG-PASS v2.0 achieved the most balanced performance across precision (98.7%), sensitivity (82.5%), and specificity (89.3%) (Table S3). DeepARG and RGI strict achieved 100% precision at a significant cost of sensitivity, particularly for ARGs with below 50% CARD identity to known ARGs (Table S2). PLM-ARG, PCM (good+fair), and ARG-PASS v2.0 (lDDT) achieved the highest sensitivity, in that order, at the cost of precision and specificity. Finally, fARGene achieved the most comparable performance to ARG-PASS v2.0 with values of 97.9%, 79.1%, and 83.3% for precision, sensitivity and specificity, respectively. Functional predictions for each candidate are presented in Table S2, performance metrics for ARG-PASS v2.0 per ARG class are presented in Table S3, and further discussion is available in the supplementary material.

### ARG-PASS v2.0 predicts novel ARGs experimentally verified from a lake metagenome

We validated ARG-PASS v2.0 on metagenomic data from Rabindra Sarobar, a man-made freshwater lake in Kolkata, India, likely receptive to anthropogenic pollutants including heavy metals and antibiotics (Ghosh et al., 2021). Candidates for protein structure prediction were identified by either an MGE co-localisation search (Table S6) or a conserved region search (CRS) (Table S7, see Methods). In total, 23 candidates were predicted functional, which confer resistance to aminoglycosides, β-lactams, or trimethoprim (Table S4). Eleven were tested in *E. coli* BW25113 (Δ*bam*BΔ*tolC*), of which ten showed activity against their respective antibiotics (Table S4, Figure 2a). A further 43 candidates were predicted functional by ARG-PASS v2.0 (lDDT) alone. Four were tested, of which one had activity (Table S5). Eight of the active candidates reached Clinical and Laboratory Standards Institute (CLSI) resistant breakpoints (CLSI, 2024a): two AAC(6′), one class A, B3 and D β-lactamase, and three dfr candidates, of which two were also resistant to trimethoprim with sulfamethoxazole (Figure 2c, Table S1). An AAC(3′), a second class D β-lactamase, and the lDDT-only AAC(2′) had activity above the negative control but below the CLSI resistant breakpoint, and therefore represent pre-resistance genes (Table S1). Notably, the pre-resistant class D β-lactamase conferred an ampicillin MIC of 4 µg/mL against a negative control of 2 µg/mL, yet was nitrocefin-positive, suggesting that a positive nitrocefin result alone is insufficient to establish clinical relevance (Figure 2b). The single false positive (APH01) contains the catalytic residues across kanamycin kinases after alignment with MAFFT (alignment S1, Thompson et al., 2002), so its inactivity may reflect limitations of heterologous expression in *E. coli*, given that the protein was identified from metagenomic data and originates from an unknown bacterial host. Functionally confirmed candidates share 48% mean CARD identity with known ARGs, but 62.4% seqID across the conserved functional regions of ARP structures identified by ARG-PASS v2.0, with a mean TM-score of 0.97 (Table S4).

**Fig 2.**
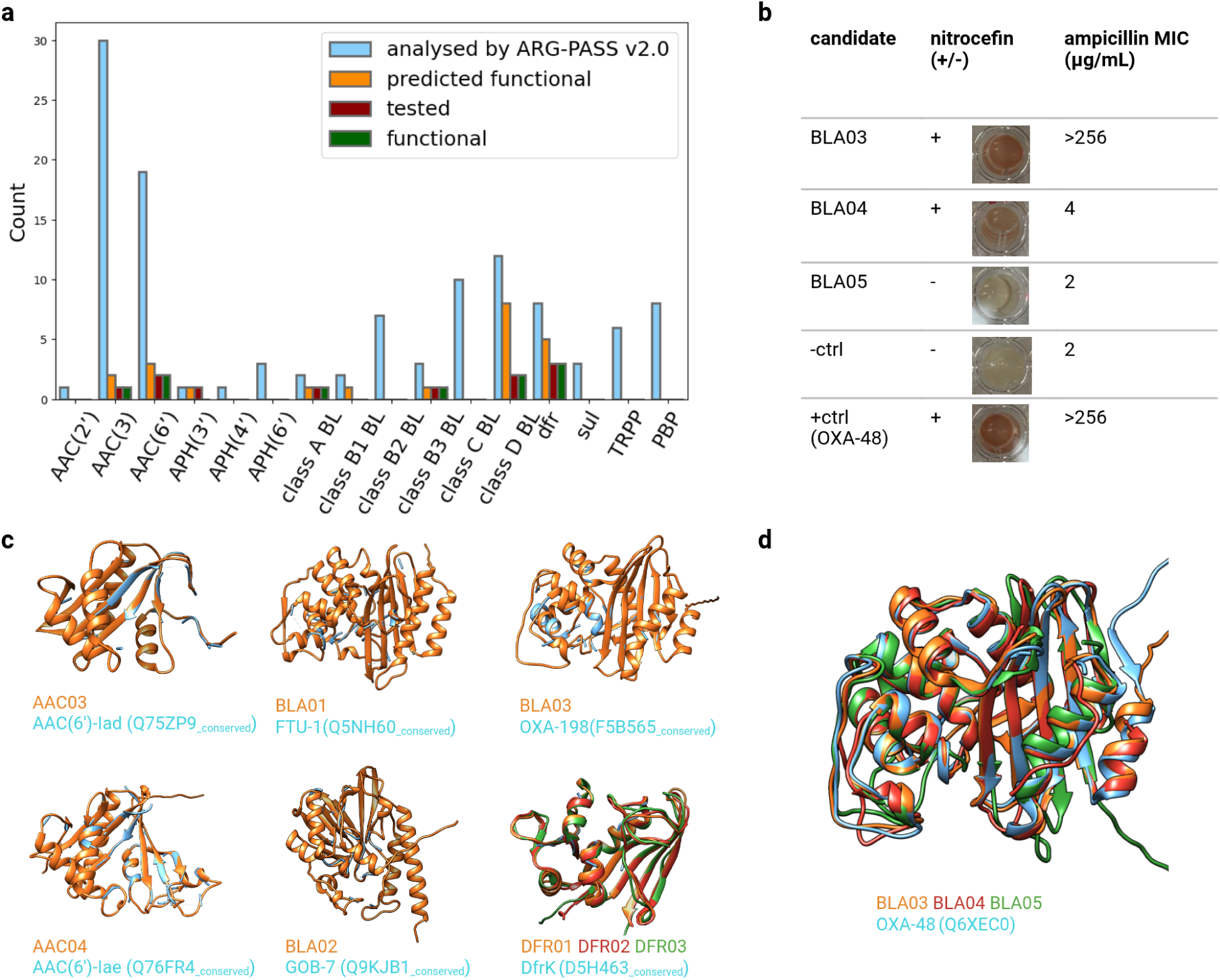
ARG-PASS v2.0 applied to an Indian lake metagenome. a, Bar chart summary of candidates analyzed, predicted functional, experimentally tested and functionally confirmed. BL: β-lactamase. b, Nitrocefin results and minimum inhibitory concentrations (MICs) for three class D β-lactamase candidates. BLA04 was nitrocefin-positive but conferred an MIC below the CLSI resistant breakpoint (4 µg/mL ampicillin). BLA05 was negative in both assays despite sharing high seqID (58%) and structural similarity (d) with known ARGs, and was correctly predicted non-functional by ARG-PASS v2.0, however misled most other tools (Table S2). c, Protein structures of candidates which conferred a clinically relevant MIC, Foldseek aligned with the conserved region of the ARP structure with which they share the highest TM-score (apart from DFR01 and DFR02). d, Protein structures of BLA03, BLA04, and BLA05 and the experimental positive control, OXA-48, aligned and visualized with USCF Chimera (Meng et al., 2006).

### ARG-PASS v2.0 predicted 918 class B1 metallo-β-lactamases across metagenomes and plasmids

Following the initial validation of our tool with the Kolkata lake samples, we wanted to apply ARG-PASS v2.0 on a larger scale. To do so, it was applied to a metagenomic database, MGnify clustered proteins (release 2024_04), and on a plasmid dataset, PLSDB (v.2024_05_31_v2).

We focused on class B1 MBLs given their role in conferring resistance to last-resort carbapenems. In total, 872 novel class B1 MBLs were predicted in MGnify (860) and the PLSDB (12, Table S8; S9). Notably, four were predicted as functional by both the class B1 and class B2 models, which each carry Asn at position 116, so they are likely class B2 enzymes (Krco et al., 2023). In addition, 46 known variants were identified (≥95% CARD identity). Most predictions were divergent from characterized enzymes: 696 (80%) share less than 60% CARD identity, with a mean of 53.2% (range 29.1–95%). Protein structures of candidates share a mean seqID of 44.2% (range 21.1–99.6%) to known class B1 MBL structures, which have an associated mean TM-score of 0.92 (range 0.74–0.99), normalized by the target structure length.

Of ten candidates predicted functional by ARG-PASS v2.0 and selected for experimental verification, nine conferred reduced β-lactam susceptibility in *E. coli* (Table S1). The tenth (MGYP003585275815) carried a mutation in the plasmid origin of replication and its expression was not confirmed. Among the nine active candidates, seven reached CLSI resistant breakpoints (five for ampicillin, meropenem and cefotaxime, and two for ampicillin alone), and two reached intermediate breakpoints against ampicillin and meropenem. The latter carries a D120E substitution in the Zn²⁺-binding site, which likely reduced the efficiency of meropenem hydrolysis. A further candidate (BLA11) was predicted functional by ARG-PASS v2.0 (lDDT) and by other tools, but not by the default mode, and showed no antimicrobial activity when tested in the lab (Table S1;S2 and Figure 1e).

Normalising to the number of MGnify studies and the number of MGnify protein clusters per biome, engineered environments (built or managed systems) yielded 368 novel MBLs (60.7 per 100 studies; 2.55 per million clusters), followed by marine 213 (14.9; 0.81), and human-associated samples 21 (2.0; 0.10). The remaining candidates occur in biomes with fewer MGnify studies. Human-associated assemblies account for more studies and a larger share of catalogue clusters than engineered environments, yet yielded roughly 4% of the engineered rate per protein cluster, which indicates that novel ARG discovery should be directed outside human-associated microbiomes.

#### Predicted marine class B1 MBLs cluster with known clinically relevant MBLs

To relate the predicted candidates to known MBLs, we constructed a maximum likelihood phylogeny of 916 MBLs, comprising all 918 candidates after dereplication at 100% identity (see Methods; Figure 3a). Known MBLs (*n* = 44) fell into two well supported clades. Clade I contained IMP, SIM-2, KHM-1, DIM-1, GIM-1, TMB-1, VAM-1, VMB-1 and PST-2 (99 tips, 25 known), and clade II contained NDM, VIM and AFM-2 (95 tips, 13 known). All were originally described from Gammaproteobacterial hosts, suggesting both clades are predominantly found in Gammaproteobacteria. Known MBLs were nested among the novel candidates rather than basal to them (median node depth 16 versus 9 in clade I, p = 1.4 × 10⁻⁷; 13 versus 8 in clade II, p = 0.003). This suggests that clinical variants emerged from within existing environmental diversity, though node depth is sensitive to sampling density.

**Fig 3.**
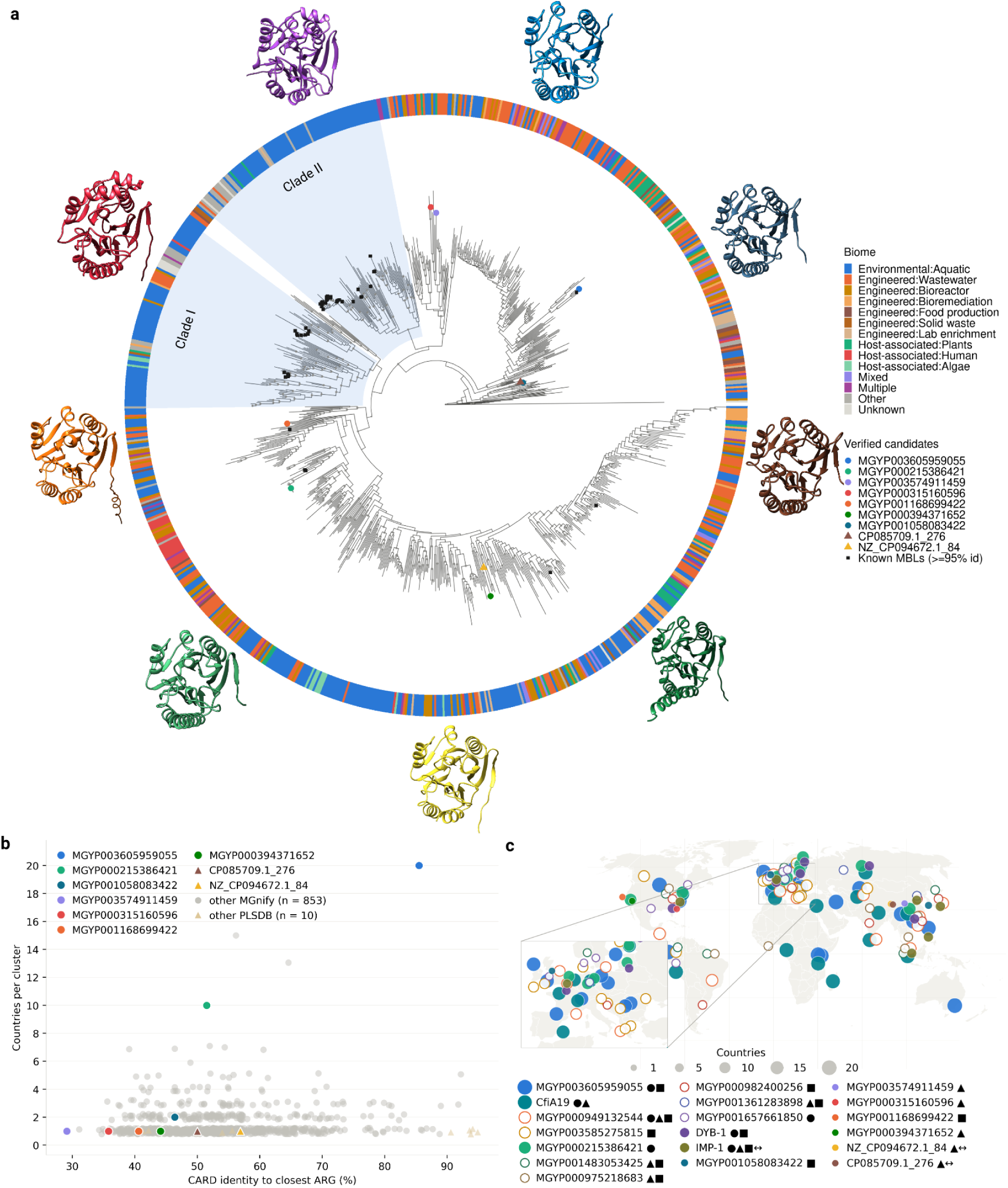
Phylogeny, sequence homology, and geographic distribution of predicted class B1 MBLs. a, Maximum likelihood phylogeny of 872 predicted and 44 known class B1 MBLs (including the 4 predicted class B2 MBLs) from MGnify and PLSDB, rooted on *E. coli* glyoxalase II. Functionally verified candidates are labelled, with their predicted structures depicted inset, visualized with USCF Chimera. b, CARD identity (%) to the nearest characterized enzymes against the number of countries occupied. Identity is that of the MGnify cluster representative and country counts include all cluster members. Functionally verified candidates are labelled and PLSDB candidates (triangles) are shown for comparison but were excluded from the correlation and group comparisons given non-comparable metadata. c, Geographic distribution of known and predicted candidates occurring in five or more countries, and of those functionally verified in this study. Functionally verified or known candidates are labelled with full circles and non-verified with empty circles in the legend. Symbols denote the biome the candidate was found: ● human-associated, ▪ engineered, ▴ environmental; and candidates with a detected MGE association are marked ↔

Marine sequences were strongly over-represented in these clades, constituting 99 of 156 novel tips (63%) compared with 115 of 716 (16%) elsewhere in the tree (OR 9.1, p = 2.9 × 10⁻³¹), as were aquaculture sequences (25 of 156 versus 23 of 716; OR 5.8, p = 2.8 × 10⁻⁸). The aquaculture signal derives from a single Chinese study (ERP108711) so should be treated as provisional, though it is not explained by country of origin, since no overall enrichment was observed among sequences originating from China (OR 1.3, p = 0.26). Independent support comes from PLSDB, where VMB-1 and VAM-1 were recovered from *Vibrio* plasmids in shrimp (Table S8) and have aquaculture metagenome sequences as their nearest relatives. MGnify candidate MGYP006226885383, sharing 42.2% seqID with VIM-29 and occurring in four countries across marine and aquaculture settings, shares 99% sequence identity with a candidate predicted from the PLSDB from *Pseudoalteromonas shioyasakiensis* (CP076682.1_588) isolated from Clayoquot Sound, Canada (Behrendt et al., 2023).

#### Class B1 MBLs in MGnify are divergent and globally widespread

To test whether sequence divergence relates to geographic distribution, we compared the range of predicted candidates across CARD identity thresholds (Figure 3b). Among the 860 novel MGnify candidates, CARD identity correlated weakly with the number of countries occupied (ρ = 0.08, p = 0.027) and not significantly with the number of biomes (ρ = 0.06, p = 0.063).

Candidates sharing less than 60% CARD identity with a known ARG occupied marginally fewer countries than those above (mean 1.41 versus 1.72, p = 0.02). However, this reflected cluster size rather than geographic range: higher-identity clusters contained more members (median 3 versus 2, p = 0.044), and the two groups had the same number of countries per cluster member (median 0.50 in both, p = 0.565). Divergent class B1 MBLs are therefore as geographically widespread as their conserved counterparts, so identity to known ARGs gives little indication of a determinant’s distribution.

#### Class B1 MBLs track their hosts’ distribution

Candidates occurring in three or more countries, as well as the functionally verified enzymes, were examined for neighbouring MGEs. The 10 open reading frames (ORFs) flanking each gene were screened against mobileOG-db (Brown et al., 2022) in up to five assemblies per cluster. Only one out of 97 novel candidates carried a conclusive MGE signature, suggesting that the widespread and functionally confirmed candidates are mostly chromosomally encoded. However, genetic context was limited in contigs (median 2.0 flanking genes per candidate; Table S11).

To further assess their risk, we cross-referenced their recorded biomes for human-association and queried if any taxonomic hosts have been reported as opportunistic pathogens (Table S11). Several are confined to human-associated biomes, so their spread tracks their hosts’ association with humans. MGYP000537499426 shares 99.2% sequence identity with DYB-1, deposited in MGnify in 2018 from a Chinese study of fermented beverages (L. Zhang et al., 2014). DYB-1 was not described until 2024, following a fatal *Indolivaga capnocytophagoides* peritonitis in a patient treated with meropenem (Imai et al., 2024). Its closest known homologs at the time were CfiA variants at approximately 50% sequence identity. *I. capnocytophagoides* is a gut anaerobe and recognized opportunistic pathogen. In our data, DYB-1 occurs in six countries and, apart from the fermented-drink biome, only in human-associated biomes including gut metagenomes from people in China (Figure 3c, Table S9; S10). Comparably, five novel and geographically widespread candidates are recorded only from human-associated biomes (Table S9). For example, MGYP000215386421 (hereafter BLA14) and MGYP001657661850 both share ∼50% identity with CfiA variants (Figure 3c, Table S9) and taxonomic analysis suggests *Alistipes* sp. and *Desulfovibrio* sp., respectively (Table S11). Notably, both were recently implicated in cases of bacteraemia in Japan (Watanabe et al., 2025, 2026). MGYP001657661850 shares 97% identity with the CfiA-like gene of *D. fairfieldensis* KML2502 (GAB7239148.1) (Watanabe et al., 2026). Clinical *A. finegoldii* isolates were reported carbapenem-susceptible (Watanabe et al., 2025), and a protein which shares 96.5% identity with BLA14 was predicted by Berglund et al. (2017) which originated from *A. finegoldii* DSM 17242 (NC_018011.1), but was negative on a Carba NP test (measuring carbapenem hydrolysis) after heterologous expression without codon optimisation. In this study, BLA14 conferred resistance exceeding the clinical breakpoint for ampicillin, meropenem, and cefotaxime (Table S1) which indicates that the previously reported lack of activity may reflect differences in experimental conditions rather than an intrinsic absence of enzymatic activity. Whether the gene is expressed in *A. finegoldii* is unknown, though a resistance phenotype could arise as described for CfiA in *Bacteroides fragilis*, where insertion elements drive expression (Lee et al., 2026).

A further eight candidates occur in human-associated biomes alongside at least one other biome type, highlighting hosts with broader ecological range (Table S11). The most widely distributed candidate, MGYP003605959055, was recovered from 20 countries across engineered and human-associated biomes and assigned to *Empedobacter* sp., a genus containing opportunistic pathogens.

### Clinically relevant ARGs in fermented soybean paste are detected in human gut samples

The occurrence of DYB-1 in both fermented beverage and human gut metagenomes from the same country raises the possibility that fermented foods provide a route by which exogenous ARGs reach the human microbiome. Screening the UHGP-95 catalogue (Almeida et al., 2021) with ARG-PASS v2.0 predicted 23 novel class B1 MBLs (Table S12), one of which (MGYP001058083422), from *Bacillus sonorensis* recovered from gut samples from China, matched a candidate recovered from a fermented soybean paste (*doenjang*) metagenome (Kim et al., 2017). This enzyme (renamed BLA15) conferred clinical levels of resistance to ampicillin, cefotaxime and meropenem (Table S1). Extending the screen to all ARG classes in the doenjang metagenome (Table S13) identified three further novel candidates in organisms recovered only from gut samples from South Korea and China (Tables S14): a class A β-lactamase and an ANT in the same *B. sonorensis* strain, and a second class A β-lactamase in *Bacillus licheniformis* strains from both countries. The doenjang and gut strains share ≥99.7% ANI (Table 1, Table S15). When expressed in *E. coli*, the two class A β-lactamase conferred clinical levels of resistance to ampicillin and cefotaxime, while the ANT conferred clinical levels of resistance to streptomycin (Table 1).

**Table 1:** Resistance profile of *Bacillus* spp. identified from doenjang and human gut samples from East Asian populations.

| Food strain (doenjang) | Gut isolate · country · ANI · MGnify accession | ARG class | Best hit (CARD identity %) | AMP | CTX | MER | STR |
| --- | --- | --- | --- | --- | --- | --- | --- |
| *Bacillus sonorensis* | AM12-57B · China · 99.92% · MGYG000000227 | Class B1 MBL | Bla2 (46%) | + | + | + |  |
| | | Class A β-lactamase | Bcl (67%) | + | + | | |
|  |  | ANT | ANT(6)-Ib (53%) |  |  |  | + |
| *Bacillus licheniformis* | CBA7132 · South Korea · 99.7% · MGYG000002357<br>AF39-10 · China · 99.68% · MGYG000002357 | Class A β-lactamase | BcIII (64%) | + | + | | |

ANI: Average nucleotide identity; AMP: ampicillin; CTX: cefotaxime; MER: meropenem; STR: streptomycin; +:conferred an MIC at or above a CLSI resistant breakpoint.

All four confirmed ARGs lie on chromosomal contigs of the *B. sonorensis* and *B. licheniformis* assemblies, with no MGE signatures within ten ORFs on either side. Both species are spore-forming generalists found across environmental and fermentation microbiomes (Palmisano et al., 2001), suggesting these determinants are distributed with their hosts rather than by LGT. The class A β-lactamase from *B. licheniformis*sits within an intact regulon encoding its own β-lactam sensor (blaR1) and repressor (BlaI/MecI/CopY family). Similar architecture is mobilized elsewhere: BLA15 shares approximately 80% identity with a plasmid-borne B1 MBL from *Bacillus* sp. CMF21 (PLSDB CP085709.1), also functionally confirmed here as BLA06 (Table S1), which is flanked by a similar blaR1–blaI module, and a near-identical gene and module occur on a separate plasmid in *Metabacillus dongyingensis* (NZ_OK052578.1). Self-regulating β-lactamases of soil-associated Bacillales are therefore capable of plasmid mobilisation, offering a possible route by which the *B. sonorensis* class B1 MBL could reach pathogenic species in the human gut. A phylogenetic tree of MGnify and doenjang candidates whose closest matches in CARD are a class B1 MBL from *Bacillus* spp., is presented in Supplementary Figure 1.

## Discussion

Sequence similarity methods primarily identify ARGs resembling those already characterized, and their effectiveness declines at low sequence identities, which is precisely where uncharacterized diversity is greatest. ARG-PASS v2.0 addresses this limitation by making predictions relying on functionally important protein structure regions. Ten of 11 candidate ARGs tested from the Kolkata lake metagenome were experimentally verified and eight reached CLSI resistant breakpoints, which establishes the method as a more precise improvement over ARG-PASS v2.0 (lDDT), as well as competing computational methods based on performance on the benchmarking dataset. This likely reflects the use of evolutionary rather than structural conservation to identify functionally important residues (see Methods). Prioritising candidates in the lake metagenome by MGE proximity, as done elsewhere (Razavi et al., 2020; A.-N. Zhang et al., 2021), recovered only six functional predictions, five of them dfr proteins, and two co-localising with thyA, which associates with dfr independently of MGEs (Domingo et al., 2011). The conserved region search (CRS) was a complementary screen and recovered a further 20 functionally predicted candidates. The CRS is readily generalisable to other protein families in large metagenomic data. However, it failed to detect the dfr and AAC(2’) candidates, likely because the conserved region of ARP structures belonging to dfr and AAC(2’) was too short to produce a significant alignment (see Supplementary Discussion). Flanking regions of potential MGEs were searched for ARGs with relaxed alignment thresholds, so any candidate identified by the CRS near a recognisable MGE would have been identified by both methods. Only one was, which indicates the remainder are potentially chromosomal.

Applying ARG-PASS v2.0 to MGnify and the PLSDB asked whether structure-based prediction can identify which novel ARGs merit surveillance. Most identified candidates are highly divergent from characterized enzymes, yet as geographically widespread. Most also lie outside human-associated environments, and genetic context analysis suggests they are also predominantly chromosomal, so their distribution likely tracks that of their hosts. Cross-referencing biome and taxonomic assignments identifies ARGs restricted to human-associated biomes, as well as those recovered across a wider ecological range. In the former, the host is in several cases a reported opportunistic pathogen. Had structure-based prediction models such as ARG-PASS been applied to MGnify before the fatal clinical case reported in 2024, DYB-1 would have been flagged years earlier. Predicted class B1 MBLs in *Alistipes finegoldii* and *Desulfovibrio* sp. follow the same pattern and warrant similar attention. Novel ARGs recovered from a doenjang metagenome, chromosomally encoded in *Bacillus sonorensis* and *B. licheniformis*, were also found in gut samples from populations consuming fermented soybean products. Both species are spore-forming and distributed across soil, plant, and animal-associated environments (Muras et al., 2021; Palmisano et al., 2001; Raza et al., 2019) and were recently found resistant to β-lactams and streptomycin from African bread starters (Adimpong et al., 2012).

All six experimentally verified ARGs identified with a human-association in this study (3 class B1 MBLs, two class A β-lactamases, and an ANT) reached CLSI ‘resistant’ breakpoints in a hypersusceptible *E.coli* background. Ruppé et al. (2019), found little evidence of ARG mobilisation between gut-associated species. However, their functional criterion differed by assessing activity against a negative control or a nitrocefin test rather than clinical breakpoints. Such criteria may overstate relevance, as a nitrocefin-positive class D β-lactamase from the lake metagenome described here conferred a pre-resistant ampicillin MIC (4 µg/mL). Enzymes too inefficient to confer a selective advantage would be less expected to mobilise and may have contributed to the result from Ruppé et al. (2019), however experimental differences cannot be excluded. Nevertheless, the ARGs described here confer resistance exceeding clinical breakpoints even in the absence of functional efflux pumps, which demonstrates efficient enzymatic activity. Moreover, several candidates originate from Gram-positive hosts such as *Bacillus* spp., yet retain activity when heterologously expressed in *E. coli*. Although their sequences were codon-optimized, these findings suggest that they may also function in other Gram-negative species. An MGE association was also not observed for these ARGs, but the plasmid-borne relatives of the *B. sonorensis* class B1 MBL shows the transition is possible within this family, and the selective conditions for it exist in East Asia where carbapenem resistance is increasing (Zha et al., 2025).

*B. sonorensis* carrying the class B1 MBL was present in gut isolate genomes from China but not South Korea. South Korean isolates are heavily underrepresented in UHGG (67 genomes against 56,733 Chinese) (Almeida et al., 2021), so the organism may be present at abundances too low to have been recovered from this collection. However, *B. licheniformis* was still recovered from gut metagenomes from South Korea, which suggests *B. licheniformis* could be more abundant in doenjang and/or better able to survive gut passage. Exposure to *B. sonorensis* may also be higher in China, where *B. sonorensis* is a known member of fermented soybean products including soy sauce (W.-F. Yin et al., 2012) and douchi (M. Yin et al., 2026). To our knowledge, this is the first report of novel, clinically relevant ARGs identified in species from fermented food with the same species recovered from populations consuming that food. Whether *B. sonorensis* and *B. licheniformis* are resident gut members or transient after doenjang consumption remains to be established.

We also found that clinically important class B1 MBL lineages phylogenetically cluster with predicted marine diversity, which aligns with Berglund et al. (2017), who proposed an environmental origin for MBLs in *Shewanella* sp., a genus spanning freshwater and marine environments. Two independent approaches now place the closest relatives of clinical MBLs in aquatic bacteria, which strengthens the case for an aquatic reservoir and potential origin. The association is also relevant to historical and future clinical emergence given aquaculture sequences were enriched among these clades, and one marine candidate is already plasmid-borne in *Pseudoalteromonas shioyasakiensis*.

Altogether, ARG-PASS v2.0 has identified novel ARGs across One Health compartments: both in species already resident in the human microbiome and in generalists that reach it from elsewhere. The bacterial host may be an opportunistic pathogen, or the gene may subsequently transfer to a pathogen. In both cases, antibiotic treatment can be compromised. ARGs which are chromosomal or whose context cannot be resolved on short contigs, demonstrate that effective resistome surveillance should attempt to resolve the host association of an ARG rather than only prioritising MGE proximity to infer risk (A.-N. Zhang et al., 2021).

Several limitations of this study should be acknowledged. Candidates were verified by heterologous expression in *E. coli,* which may not reflect function in native hosts, and a larger portion of sequences used in benchmarking ARG-PASS v2.0 against existing computation tools were functional (*n* = 280) as opposed to non-functional (*n* = 27), which can inflate precision metrics across tools. Experimental verification remains the principal bottleneck in ARG discovery, so although structure-based prediction would have identified DYB-1 before its clinical description, verification would still have been likely required before inclusion in databases such as the CARD. Inference of chromosomal location for MGnify candidate was constrained by contig length, with a median of 2 flanking genes available against a possible 20, so some candidates may have undetected MGE associations. ARG diversity here is relative to the CARD, which is populated predominantly by clinically derived sequences, thus recovery rates across biomes partly reflect prior characterisation as well as underlying diversity. Finally, machine learning requires existing examples and cannot identify ARGs unlike those already described, so we stress the importance of functional metagenomics and other computational methods for discovering completely new resistance mechanisms.

When applied to existing sequence data, ARG-PASS v2.0 can link previously undetected ARGs to their hosts and environments. Looking forward, we envision its deployment on sequence data generated during molecular diagnostics (particularly where organisms are unculturable or no homologs exist in ARG databases), altogether informing antibiotic therapy and stewardship in the clinic.

## Methods

Unless stated otherwise, all steps follow ARG-PASS as described previously (Bartrop et al., 2026). Below we describe the components that differ in ARG-PASS v2.0, together with the screening and analysis performed here.

### Measures of sequence identity

Two measures of amino acid sequence identity are reported and are not interchangeable. “SeqID” refers to identity over the aligned regions of two protein structures, taken from Foldseek structural alignments in TM-align mode. “CARD identity” refers to sequence identity between a candidate and its closest match in the custom CARD database of the ARGs used to train the one-class SVM models, computed from Diamond sequence alignments. Because structural and sequence alignment tools pair residues differently, the two measures can differ by several percent.

### ARG-PASS v2.0 (default and lDDT)

#### Training set

The training set of 1,193 ARP structures encoded by well-characterized ARG classes in CARD v4.0.1 (Alcock et al., 2023) (aminoglycoside acetyltransferases (AAC), aminoglycoside nucleotidyltransferases (ANT), aminoglycoside phosphotransferases (APH), class A, B, C, and D β-lactamases, dihydrofolate reductases (dfr), mobile sulfonamide-resistant dihydropteroate synthetases (SUL), and tetracycline-resistance ribosomal protection proteins (TRPP)) were obtained from the AlphaFold database (AFDB) or predicted with AlphaFold2 (AF2, Jumper et al., 2021) or AlphaFold3 (AF3, Abramson et al., 2024). β-lactam-resistant penicillin binding proteins (PBPs) were not used in training ARG-PASS v2.0 (default), however remain in ARG-PASS v2.0 (lDDT), as resistance is determined by distinct point-mutations which are not expected to be captured as conserved residues across PBPs by CONSTRUCT (see below).

For all β-lactamase classes (except class B2), redundancy was reduced by clustering at 99% seqID over 90% length using Foldseek v.10-941cd33 (van Kempen et al., 2024). Structures were then sorted by class and clustered at 20% seqID, with clusters containing fewer than four structures excluded in ARG-PASS v2.0 (default), and fewer than three structures in ARG-PASS v2.0 (lDDT). AlphaFold/UniProt accessions for ARGs used in training are available in Table S16.

#### Conserved region identification - ARG-PASS v2.0

The principal advance of ARG-PASS v2.0 over the original ARG-PASS method (Bartrop et al., 2026) is more precise identification of functionally important amino acid residues within ARP structures, which refines the seqID vs TM-score distributions used for functional prediction. Conserved regions were identified with CONSTRUCT (Chivot et al., 2025). CONSTRUCT addresses a limitation of Rate4Site (Pupko et al., 2002): rather than treating site-specific substitution rates as independent, CONSTRUCT weights each site’s evolutionary rate by those of its spatially proximate neighbours, determined from Euclidean distances between residues in a representative structure. The representative structure used in all cases had a predicted Local Distance Difference Test (pLDDT) > 90. Patches of conserved sites likely to be functionally critical are thereby identified from spatially correlated site-specific substitution rates, retained at a threshold of log(p) ≥ 8 (default setting).

For each 20% seqID cluster, ARP structures were split into training and held-out sets, depending on the number of ARP structures within the cluster (Table 2). A multiple sequence alignment of the amino acid sequences within each respective training set was then generated with MAFFT v.7.526 (Katoh et al., 2009`;--maxiterate 1000 --localpair`) required as CONSTRUCT input, and it then ran under default settings. This was repeated progressively with the CONSTRUCT threshold relaxed each time in increments of 0.05, starting from a stringent starting value, which increased the number of residues extracted and thus relaxed the stringency of the model. For each CONSTRUCT threshold, the model was trained with a one-class support vector machine (SVM, see below), and the first threshold satisfying the criterion for the held-out set was selected (Table 2). A MAFFT alignment was then regenerated for all amino acid sequences in the cluster and residues were extracted from each ARP structure using the selected CONSTRUCT threshold. Notably, clusters containing fewer than four ARP structures were omitted. This is because the one-class SVM models require at least three training points to generate a reliable DB, meaning leave-one-out cross-validation requires a minimum of four ARP structures.

**Table 2.** Training scheme for CONSTRUCT threshold selection.

| Number of ARP structures in cluster | Training scheme | Selection criterion |
| --- | --- | --- |
| 4–10 | Leave-one-out cross-validation | $\geq 50\%$ of held-out set predicted functional across all folds |
| 11–20 | 5-fold cross-validation | 1 predicted non-functional from |
|  |  | held-out set across all folds |
| 21–100 | 5-fold cross-validation | 100% of held-out set predicted functional across all folds |
| >100 | Single 80:20 train:held-out split | 100% of held-out set predicted functional |

#### Conserved region identification - ARG-PASS v2.0 (lDDT)

Conserved regions were identified and extracted in ARG-PASS v2.0 (lDDT) based on structural conservation (Bartrop et al., 2026; Gilchrist et al., 2026). For each 20% seqID cluster of ARP structures, structurally conserved residues were extracted using the median, 75th percentile, and 90th percentile of the distribution of lDDT scores across the structural alignment. However, in this case, ARG-PASS v2.0 (lDDT) differs from the original ARG-PASS method by the use of the training data described here and candidate assignment to only 20% seqID clusters for functional prediction. Structurally conserved residues were extracted across both class B1 and B2 MBL structures without de-replication at 99% seqID in ARG-PASS v2.0 (lDDT), thus combining their prediction into one model.

The increased sensitivity of ARG-PASS v2.0 (lDDT) compared to the default configuration likely derives from using structural rather than evolutionary conservation to identify functionally important regions. Structurally conserved regions yield high lDDT scores across structurally aligned regions, however they don’t necessarily map to functional conservation. Protein backbones often tolerate more substitutions than catalytic sites, exemplified where similar structures can have different functions (Koehler Leman et al., 2023). The extracted residues are therefore less constrained at the amino acid level, widening the range of sequence identities in the resulting seqID vs TM-score distribution. Functional prediction proceeds by aligning these regions to the candidate, so a protein sharing the structurally conserved core will be more likely predicted functional even where the catalytic residues differ. This likely accounts for the mode’s greater sensitivity yet lower precision, as illustrated by DFR04, BLA05, and BLA11, which were predicted functional by ARG-PASS v2.0 (lDDT) but showed no activity in this study, and AAC01, also predicted functional by ARG-PASS v2.0 (lDDT) but not the default mode, and had minor resistance activity (Table S1).

#### Training and one-class SVM hyperparameter selection

Training data comprises all-versus-all pairwise comparisons of conserved regions of ARP structures within each 20% seqID cluster, to produce an *x*, *y* distribution of seqIDs (*x*) vs TM-scores (*y*). Comparisons were computed with Foldseek TM-align (`--alignment-type 1--exhaustive-search`), with TM-scores normalised by subject length, and distributions standardised using scikit-learn’s `StandardScaler` (Pedregosa et al., 2011). For ARG-PASS v2.0 only, pairs in which the query was not 90–110% of the subject length were excluded from clusters containing more than 10 structures. Normalising by subject length depresses TM-scores for shorter queries and inflates them for longer ones, so this filter prioritises precision at a slight cost to sensitivity.

One-class SVMs were trained on each distribution using an RBF kernel with ν = 1/N, where N is the number of training points, and γ set by scikit-learn’s `scale` option. Setting ν = 1/N assumes all training data are inliers, which avoids underfitting and limits bias from clusters containing many near-identical variants such as TEM β-lactamases.

### Functional prediction

In the original ARG-PASS method, candidates were assigned to structure cluster levels (20%, 30%, or 50%) for functional prediction according to their existing homology to a known ARP structure. However, in both configurations of ARG-PASS v2.0 (default and lDDT), ARP structures are clustered at 20% seqID only, so all candidates are aligned to conserved regions of ARP structures at this single cluster level.

Each candidate structure is aligned to the Foldseek databases of conserved regions of ARP structures, to obtain an *x*, *y* distribution of seqIDs (*x*) vs TM-scores (*y*) for the candidate structure (also scaled using scikit-learn’s ‘StandardScaler’). As with the training data, comparisons were computed with Foldseek TM-align (`--alignment-type 1--exhaustive-search`), with TM-scores normalised by the conserved region of the ARP structure (subject) length.

Due to the alignment of short, conserved regions of ARP structures against full-length candidate structures using Foldseek TM-align, seqID calculations were underestimated due to alignment gaps not being properly accounted for in the aligned region of the candidate structure. Manual adjustment of seqIDs was required by incorporating an alignment length of the conserved region of the ARP structure (subject length).

### Benchmarking against existing computational tools

Both configurations of ARG-PASS v2.0 (default and lDDT) were benchmarked against existing computational tools designed for ARG prediction, through the use of strictly experimentally verified positive (*n* = 280) and negative (*n* = 27) candidate ARGs, across various CARD identities (22.5 - 99.2%). A candidate was defined as positive if it conferred an MIC above that of a negative control in *E. coli*. It was confirmed that no positive candidates are in the CARD, which ensured a fair comparison between computational methods. Candidates were compiled from various sources: ARGs identified by pairwise comparative modelling (PCM) applied to the MetaHIT catalogue (Ruppé et al., 2019); ARGs curated in ResFinderFG v2.0 from functional metagenomics (Gschwind et al., 2023); ARGs identified by the original ARG-PASS method from Human Microbiome Project (HMP) reference strains, including negative predictions confirmed non-functional after testing (Bartrop et al., 2026); and candidates experimentally tested in this study. Compilation of the PCM and ResFinderFG v2.0 candidates, including sequence retrieval, deduplication, and class-balanced sampling, followed the procedure described previously (Bartrop et al., 2026).

Comparator computational tools were DeepARG v2.0.0 (Arango-Argoty et al., 2018), PLM-ARG (Wu et al., 2023), fARGene v0.1 (Berglund et al., 2019), PCM v1.0 (image pcm_202010.img) (Ruppé et al., 2019), and the CARD Resistance Gene Identifier (RGI) (Alcock et al., 2023). PLM-ARG predictions were called at a probability threshold of 0.5 (default), any hit was considered for DeepARG and fARGene, only “strict” hits for CARD RGI were retained, and PCM hits were considered which fell into “good” and “good and fair”. All tools were run under default settings unless specified. Benchmarking metrics were calculated as follows: precision = TP/(TP+ FP), sensitivity = TP/(TP + FN) and specificity = TN/(TN + FP). Candidates for which a tool returned no prediction were excluded from that tool’s calculation: ARG-PASS v2.0 does not predict PBPs, fARGene does not predict ANT, PBPs, dfr, and SUL, PCM does not predict PBPs, and deepARG does not predict SUL.

### Datasets and candidate identification

#### Conserved Region Search (CRS)

To enable high-throughput screening of large sequence datasets for likely ARG candidates, we introduce the Conserved Region Search (CRS). CDS were queried with Diamond v0.14.152 (Buchfink et al., 2015) against a database comprising only the conserved regions of ARP structures identified by CONSTRUCT during ARG-PASS v2.0 development, with the arguments`--ultra-sensitive --max-target-seqs 1`. It was imperative that gaps remained intact in the place of non-conserved residues in the MAFFT alignment otherwise a decrease in sensitivity and more spurious alignments were observed. We applied the CRS screen to the benchmarking dataset and achieved a recall of 81% of functional sequences (Table S2, see Supplementary discussion).

#### Lake metagenome

Raw reads for the lake metagenome samples (Rabindra Sarovar in south Kolkata, India) were obtained from the NCBI SRA database (SRA accessions SRR8209851, rainy season; SRR8209852, winter season) (Ghosh et al., 2021), adapter-trimmed using Fastp v0.23.2 (Chen et al., 2018), assembled with Megahit v1.2.9 (D. Li et al., 2015), and protein-coding sequences (CDS) were predicted using Prodigal v.2.6.3 (Hyatt et al., 2010).

Candidates for structure prediction were identified by two methods. First, potential mobile genetic elements (MGEs) were identified by aligning CDS with Diamond against mobileOG-DB (Brown et al., 2022;`--id 80 --query-cover 80 --subject-cover 80`), and by HMMER v3.1 search against PFAM (Mistry et al., 2021; *E*-value ≤ 0.001), retaining annotated hits matching the keyword strings “DDE”, “Transpos”, “Resolvase”, “Replicase”, “Tnp”, “Integr”, “Phage”, and IS family identifiers. CDS within 10 ORFs of a putative MGE were then screened for ARGs by Diamond alignment to a custom Diamond database of full-length sequences of ARGs obtained from the CARD, which include those used to train ARG-PASS v2.0 (--ultra-sensitive --query-cover 50 --subject-cover 50), with results presented in Table S6. Second, CDS were screened by the CRS as described above (Table S7), however ARGs belonging to TRPP, SUL, and PBP were omitted for this screen as they returned more than 200 hits each, likely because the conserved regions of their ARP structures are longer than other classes, as specified in the training scheme (Table 2).

#### PLSDB, MGnify, UHGP-95, and Doenjang

The 90% clustered MGnify protein database (2024_04) (Mitchell et al., 2020) and UHGP-95 v3.0 (Almeida et al., 2021) were downloaded from the MGnify FTP site, and CDS for the doenjang metagenome (ERP111597) directly from MGnify. PLSDB v2024_05_31_v2 (Molano et al., 2025) sequences were downloaded from the host website and CDS predicted with Prodigal.

CRS was applied to each dataset as described above targeting class B1 MBLs only, whereas all ARG classes were targeted for the doenjang metagenome (Table 3). CRS sequence hits from all datasets apart from UHGP-95 were clustered at 95% identity with CD-HIT over any length (W. Li & Godzik, 2006). Doenjang candidates were dereplicated on this basis, since all derive from a single metagenome. For PLSDB and MGnify, each novel candidate carries its own geographic and biome record, so all were retained. The PLSDB candidates were non-redundant apart from one pair sharing 99.6% identity. Ten MGnify candidates, including the functionally verified MGYP000215386421, fell within a single group despite all being singletons in the MGnify protein database. Notably on Figure 3c, MGYP000148303205 was omitted given it shares 99.6% to MGYP000215386421 and also met the 5 or more country distribution criteria.

**Table 3.** Overview of candidates identified and predicted functional by ARG-PASS v2.0 from databases analyzed in this study.

| Database | Version/study | ARG classes targeted | Pre-screening | Candidate structures predicted | Structure Prediction tool | Candidates predicted functional | Selected for experimental verification · tested |
| --- | --- | --- | --- | --- | --- | --- | --- |
| Lake metagenome | mgm4875225.3 (rainy);<br>mgm4875226.3 (winter) | all | MGE association;<br>CRS | 116 | AF2; AF3 | 23 | 12 · 11 |
| PLSDB | v.2024_05_31_v2 | class B1<br>MBL | CRS | 15 | AF3 | 12 | 2 · 2 |
| MGnify 90% protein clusters | 2024_04 | class B1<br>MBL | CRS | 1014 | ColabFold | 860 | 9 · 8<sup>^</sup> |
| UHGP-95 | v3.0 | class B1<br>MBL | CRS | 26 | AF3 | 23 | 1 · 1<sup>^</sup> |
| Doenjang metagenome | ERP111597 | all | CRS | 70 | AF3 | 4 | 4 · 4<sup>^</sup> |

<sup>^</sup>: one identical candidate (class B1 MBL) exists across MGnify, UHGP-95, and doenjang

#### Candidate filtering and protein structure prediction

CRS sequence hits from all datasets were aligned to the custom Diamond database of full-length ARG sequences from the CARD (`--ultra-sensitive --max-target-seqs 1 --query-cover 80 --subject-cover 80 -e 10 --id 0`). Sequences were discarded if they matched a known ARG at ≥95% identity, did not begin with a standard AUG start codon, or fell outside 70–130% of the subject length. Putative class B3 MBLs were removed from the MGnify candidate set on the basis of the Zn²⁺-binding motif (HXHXDH), which is distinct from that of class B1 MBLs (Krco et al., 2023).

Protein structures encoded by amino acid sequences from the CRS and/or the MGE co-localisation search identified from the lake metagenome, PLSDB, UHGP-95, and the doenjang metagenome were predicted with AF2 or the AF3 webserver as previously described (Bartrop et al., 2026). Due to the larger number of MGnify candidates (*n* = 1014), these were folded with ColabFold v1.6.3 (Mirdita et al., 2022) on a Digital Research Alliance of Canada cluster. All predicted structures were then functionally classified by ARG-PASS v2.0 (lake metagenome: Table S1; PLSDB: Table S8; MGnify: Table S9; UHGP-95: Table S12; doenjang metagenome: Table S13). For comparison between ARG-PASS v2.0 sensitivity modes, candidates identified from the lake metagenome (Table S5), MGnify, and the doenjang metagenome (data not shown) were also functionally classified by ARG-PASS v2.0 (lDDT). Per-dataset candidate counts and outcomes are summarized in Table 3.

#### Congruence of ARG-PASS v2.0 functional predictions from MGnify between ColabFold and AF3 structures

To assess whether functional predictions depend on the computational structure prediction tool, we re-folded a subset of MGnify candidates with AF3 and compared ARG-PASS v2.0 classifications to those from ColabFold structures (Table S17). All positive candidates with ≤35% identity to their closest CARD match were included, together with five randomly selected from each of three higher-identity bins (35–50%, 50–75%, 75–100%). We also re-folded ten negative candidates, five each from above and below 50% identity. Three candidates classified as functional from ColabFold structures were non-functional from AF3, all with <35% identity; which were reclassified as non-functional. Classifications were concordant for the 15 positive candidates above 35% identity and for all ten negatives, suggesting that discordance is only confined to the lowest-identity range.

### Phylogenetic analyses

Predicted class B1 MBLs from MGnify and PLSDB were combined with characterized enzymes identified by the same screen, and glyoxalase II from *E. coli* (GLO2_ECOLI, P0AC84.1) was included as an outgroup (Figure 3a). Sequences were dereplicated with CD-HIT at 100% identity over full length, aligned with MAFFT v7.526 (--maxiterate 1000 --localpair), and the alignment trimmed with trimAl v1.5.1 (Capella-Gutiérrez et al., 2009) using -gappyout. Two PLSDB entries, NZ_MZ581331.1_32 and NZ_OP958859.1_178 (encoding IMP-13 and VAM-1, respectively), were identical to MGnify candidates and are represented by those tips, giving 916 sequences (872 MGnify, 44 PLSDB) plus the outgroup. A maximum-likelihood tree was inferred with IQ-TREE 3.0.0 under the model Q.PFAM+R10, selected by ModelFinder under BIC, with 1000 ultrafast bootstrap replicates and 1000 SH-aLRT replicates. The tree was rooted on the outgroup. Nodes were considered well supported at SH-aLRT ≥ 80 and ultrafast bootstrap ≥ 95; 448 of 916 internal nodes (49%) met both criteria. Candidates sharing ≥95% CARD identity (*n* = 44) were designated as known and the remainder as novel. The tree was visualized with ggtree and ggtreeExtra in R v4.5.2.

For candidate class B1 MBLs from *Bacillus* spp., those predicted from MGnify, PLSDB, and the doenjang metagenome which share a closest homolog with a class B1 MBL originating in *Bacillus* spp. were aligned with MAFFT as above, together with the characterized enzymes BcII (*Bacillus cereus*, AAA22562.1) and Bla2 (*Bacillus anthracis*, AAR20596.1). A maximum-likelihood phylogeny was inferred with IQ-TREE 3.0.0 (`-m MFP -B 1000 -alrt 1000 -T AUTO`), rooted on the glyoxalase II outgroup (GLO2_ECOLI, P0AC84.1). LG+I+G4 was selected by ModelFinder under BIC, and node support was assessed by 1,000 ultrafast bootstrap replicates and 1,000 SH-aLRT replicates (Figure S1). The final tree was visualized with FigTree v.1.4.4 (available http://tree.bio.ed.ac.uk/software/figtree/).

### ANI calculation

Four metagenomic assemblies from the doenjang study (ERZ776998, ERZ777000, ERZ777001, ERZ777002) were retrieved from MGnify. Reference genomes were obtained from NCBI for the gut-derived isolates *B. licheniformis* CBA7132 (Korea; NZ_CP021970.1), *B. licheniformis* AF39-10 (China; QROJ01, GCF_003474315.1) and *B. sonorensis* PMC204 (China; NZ_CP139190.1). Because each assembly contained both Bacillus species alongside other community members, contigs were first assigned to species before Average Nucleotide Identity (ANI) was calculated. Assemblies were filtered to contigs ≥5 kb using seqkit, and each contig was compared individually against the three reference genomes using skani (Shaw & Yu, 2023) (dist --qi -s 70). Each contig was assigned to the reference giving the highest ANI, and contigs with ≥97% ANI and ≥80% query alignment fraction were retained. Retained contigs were pooled by species to form one *B. licheniformis* and one *B. sonorensis* bin per assembly. ANI between each bin and each reference genome was then calculated with skani (dist -s 70). skani applied its learned-ANI adjustment, a regression model trained on metagenome-assembled genomes, to account for fragmented input. Reported ANI values include the corresponding reference alignment fraction, which indicates the proportion of the reference genome represented in the comparison. Cross-species comparisons gave 84.3–84.5% ANI, confirming that the two populations were correctly separated. Per-assembly values are given in Table S15 and Table 1 in the main text reports the comparison with the highest reference alignment fraction for each species.

### Metadata of class B1 MBL candidates

#### MGnify candidates and cluster representatives

Metadata for all MGnify class B1 MBL candidates were retrieved from the MGnify protein database (release 2024_04), hosted as a Google Cloud BigQuery public dataset (`bigquery-public-data.ebi_mgnify`). Candidate accessions were joined through the `metadata`, `assembly`, `study` and `biome` tables to obtain the ENA assembly accession, ENA study accession and GOLD biome lineage for each protein. Each candidate’s best CARD hit, together with its biome, assembly and study accessions and cluster summary counts, is given in Table S9.

Cluster membership was taken from the `cluster_representative` field of the `protein` table, which for a representative equals its own accession. For each candidate, all proteins assigned to the same cluster were retrieved and joined through the same tables, so that biome, assembly and study provenance were obtained per member rather than inherited from the representative. Per-member geography, biome, and study occurrence are given in Table S10.

Geographic metadata were retrieved from ENA, with one request per unique assembly accession, by parsing `SAMPLE_ATTRIBUTE` records from the ENA browser XML API. Tag matching accommodated the varied naming conventions used by submitting consortia, including combined `lat_lon` values with hemisphere letters and coordinates stated only in free-text sample descriptions. Where these failed, the ENA Portal API was queried as a fallback.

Placeholder values (for example `99999`, `not collected`, `not applicable`) were discarded. Country was taken from the same sample record, accepting marine region or sea designations where no country was recorded, since open-ocean samples have no national attribution. Region suffixes were removed, so that entries of the form `Country: region` resolved to the country alone. Counts of countries occupied by a cluster therefore include ocean and sea regions as distinct geographic units.

#### PLSDB

Plasmid accessions for which class B1 MBL candidates were predicted in the PLSDB were queried against NCBI with EDirect, linking each nucleotide record to its BioSample to retrieve host organism, strain, isolation source, collection date, geographic location, and coordinates. Placeholder values (`na`, `missing`, `not applicable`, `not collected`, `unknown`) were treated as missing values. Geographic location strings were split on the colon into country and region, and combined `lat_lon` values were parsed to decimal degrees with hemisphere letters converted to sign.

### Signatures of mobility and taxonomy

To assess mobility, we examined the genetic context of MGnify candidates occurring in three or more countries or functionally verified in this study, together with candidates from the doenjang metagenome. For each candidate, up to 10 ORFs either side were extracted from the assembly in which it was identified. For MGnify candidates, up to five assemblies per cluster were examined given each assembly represents the genetic context of a different cluster member. Flanking proteins were aligned to mobileOG-db with DIAMOND `(--id 80 --evalue 0.001 --query-cover 80 --subject-cover 80`). Results for the MGE search for MGnify candidates are reported in Table S11.

The same flanking proteins from MGnify were assigned taxonomy with MMseqs2 (release 18-8cc5c) (Mirdita et al., 2021) against UniRef50 (Suzek et al., 2015) (`--tax-fiufieage 1 -s 6`). Assignments were restricted to Bacteria and Archaea. A consensus taxon was called per candidate at the finest rank for which at least 2 flanking proteins agreed and that taxon accounted for at least 50% of the proteins classified at that rank. The number of supporting proteins and contributing assemblies is reported alongside each assignment. In addition, candidate protein sequences occurring in 5 or more countries were searched against NCBI nr (September 2026) with the best hit by *E*-value reported. Results for the taxonomic assignment are presented in Table S11.

### Experimental verification

Candidates were selected for experimental verification from the lake metagenome to ensure representation across ARG classes. All candidates were tested which only had one predicted from an ARG class; for those with two or more (AAC(6’), class D β-lactamase, and dfr) a larger number was selected which roughly scaled with the number predicted from that class. Notably, k127_888183_4 (a predicted class B1 MBL) was unsuccessfully assembled after multiple attempts into the pGDP2 vector, and suggests toxicity in *E. coli*. For the MGnify and PLSDB candidates, MGYP001168699422 was selected due a D120E substitution in the Zn²⁺-binding site, MGYP003574911459 as it has the CARD identity to known MBLs (29.1%), MGYP001058083422 as it was also identified in the UHGP-95, and CP085709.1_276 due to its high homology to MGYP001058083422. Remaining candidates were selected to capture phylogenetic and geographic diversity. However, of these, MGYP003585275815 had no activity but contained a mutation in the plasmid ORI, and successful expression was not confirmed. For the doenjang metagenome, all candidates were selected which were predicted functional from *B. licheniformis* and *B. sonorensis,* as these strains were recovered in gut samples from East Asian populations only. Finally, seven candidates predicted functional by ARG-PASS v2.0 (lDDT) only and originating from the lake metagenome, MGnify, and doenjang datasets were tested to compare between the algorithm’s sensitivity modes.

Selected candidates and positive controls were assembled into the Minimal Antibiotic Resistance Platform (Cox et al., 2017). Codon-optimised gBlocks (IDT) were assembled into pGDP vectors by NEBuilder HiFi DNA Assembly (New England Biolabs), transformed into *E.coli* MM294, and plasmids verified by Oxford Nanopore sequencing (MinION Flongle), before electroporation into *E. coli* BW25113 Δ*bamB*Δ*tolC*. Positive controls not already present in the platform were synthesized alongside the candidates: *dfrK* (CBL80435.1) and *sul3* (ACJ63260.1), and *folA* from *E. coli* MG1655 (NP_414590.1) was used as a control to compare trimethoprim resistance from overexpression of chromosomal *folA*. No positive control was used for ANT. The Minimal Antibiotic Resistance Platform was a generous gift from Prof. Gerard Wright, McMaster University, and is available as Addgene kit #1000000143.

MICs were determined by broth microdilution (BMD) in accordance with CLSI M07 (CLSI, 2024b). Overnight MHII cultures were serially diluted and applied in duplicate to BMD plates containing antibiotic concentrations spanning the CLSI resistant breakpoint (CLSI, 2024). After 18 h at 37 °C, the MIC was recorded as the lowest concentration with no visible growth. Every plate included *E. coli* BW25113 (Δ*bamB*Δ*tolC*) carrying an empty pGDP vector as a negative control and a pGDP vector carrying a known ARG of the same class, as a positive control. A candidate was considered functional if its MIC was at least two-fold higher than the negative control in both replicates. Candidate class D β-lactamases identified from the lake metagenome were also subject to a chromogenic test (nitrocefin) for comparison to MICs by BMD. Any candidates across all databases with no detected activity by BMD were also tested on solid media. Here, 2 µL of overnight culture grown for 18h at 37°C in 5mL of LB was spotted in triplicate onto MHII agar or MHII agarose (for TMP/SMX) and concentrations of appropriate antibiotics at CLSI breakpoints. After 18h at 37°C, plates were visually inspected, with an MIC recorded as the concentration for which no visible growth was present. A candidate was considered functional if its MIC was higher than that of the negative control.

### Statistical analyses

Analyses were performed in Python 3.12 (SciPy v1.18.1 and pandas v3.0.6). All tests were two-sided, and no correction was applied for multiple comparisons. The unit of observation throughout was the cluster representative, so each distinct sequence contributes once regardless of how many times it was sampled. Studies were not used as the unit of a contingency test, since a single study contributes many proteins whose observations are not independent. PLSDB entries were excluded from biome-normalized comparisons. Enrichment of marine and aquaculture sequences within clades was assessed by Fisher’s exact test, with odds ratios reported as the sample odds ratio (*ad/bc*) from the 2 × 2 table of clade membership against biome, as returned by `scipy.stats.fisher_exact` alongside the exact *p*-value.

The same test was applied to sequences grouped by country of origin, to assess whether biome enrichment was confounded by geographic sampling. Recovery rates per biome were expressed both per 100 MGnify studies and per million protein clusters. Group comparisons (i.e., pairwise seqID and TM-score for full-length versus conserved regions of ARP structures, node depth for known versus novel tips, and the numbers of countries, biomes and cluster members for candidates above and below 60% CARD identity) used the Mann–Whitney U test. Relationships between CARD identity and the numbers of countries and biomes occupied were assessed by Spearman’s rank correlation. For the maximum-likelihood tree of MGnify and PLSDB candidates, node depth was calculated as the number of internal nodes between a tip and the root. This measure is sensitive to sampling density, and depth comparisons are interpreted accordingly.

## Supporting information

Lake metagenome ARG candidates

MGnify candidate class B1 MBLs

UHGP-95 candidate class B1 MBLs

Doenjang ARG candidates

PLSDB candidate class B1 MBLs

alignment S1

## Declarations

### Ethics approval and consent to participate

Not applicable.

### Availability of data and material

Foldseek databases of conserved regions of ARP structures and training data are available on the GitHub page. Protein sequences of candidates are available in the supplementary material. ARG-PASS v2.0 is freely available to use from the GitHub page https://github.com/btroppo/ARG-PASS.

### Authors’ contributions

LB and LPH conceived and designed the study. LB performed PCR reactions, cloning assemblies, transformations, and functional verification of novel genes. FG performed Nanopore sequencing, provided experimental guidance, and manuscript review. LPH, SR, and JB contributed to conceptual analysis, provided experimental guidance, and manuscript review. LB designed the computational method, performed bioinformatic analyses, and drafted the manuscript. All authors read and approved the final manuscript.

## Acknowledgements

This research was enabled in part by computational support provided by Calcul Québec (calculquebec.ca) and the Digital Research Alliance of Canada (alliancecan.ca).

## Supplementary Figure and Discussion

**Figure S1.**
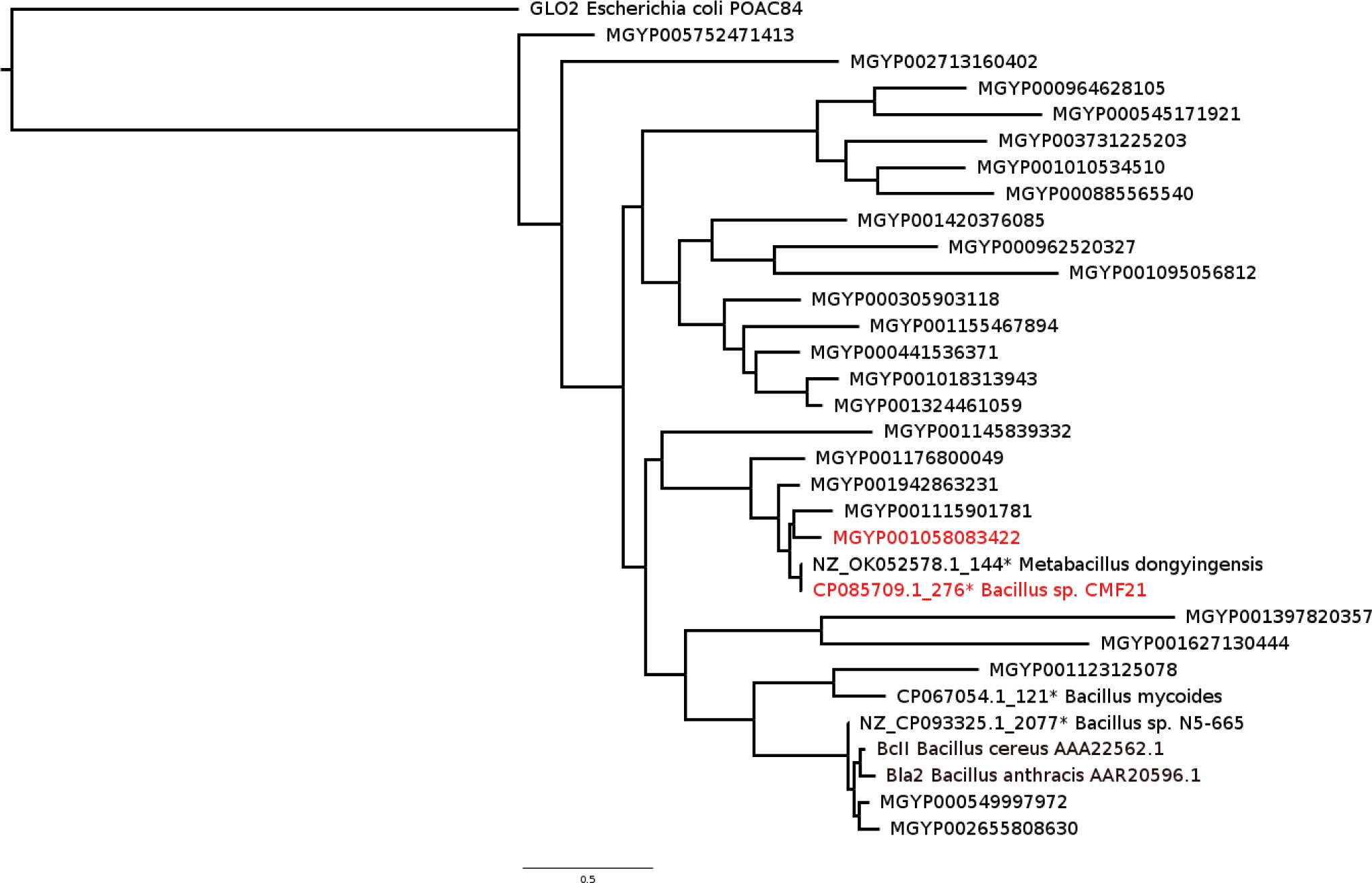
Maximum-likelihood phylogeny of known and predicted class B1 MBLs from *Bacillus* spp., rooted on the glyoxalase II outgroup (GLO2_ECOLI, P0AC84.1). Functionally verified candidates are depicted in red. *=candidates share 99.6% identity.

### Taxonomic assignment of functional ARGs identified from the lake metagenome

Two functionally confirmed candidates were placed in medium-quality metagenome-assembled genomes (MAGs) from the study which generated the sequence data (Ghosh et al., 2021).

AAC04, closest to AAC(6’)-I33 (42.9% identity), was assigned to Kapabacteriales, and BLA04, closest to OXA-29 (41.0%), to Parachlamydiales. BLA04 lay five ORFs from a Pfam phage holin domain (Phage_holin_3_6), which suggests a possible prophage context (Table S6). Notably, OXA-29 was described from *Legionella* (*Fluoribacter*) *gormanii*, an amoeba-associated freshwater bacterium occupying the same niche as Parachlamydiales, which suggests these class D beta-lactamases may be characteristic of intracellular aquatic lineages (Franceschini et al., 2001).

### Benchmarking

A number of candidates misled other tools but were correctly predicted by ARG-PASS v2.0. For example, a class B1 MBL candidate, MGnify protein MGYP001599396209 (BLA11), lacked function when expressed in *E.coli* in this study. Only deepARG and CARD strictly predicted it as non-functional. BLA11 has a best CARD identity with VIM-52 of 38.8% and contains a partial Zn2+ binding site, lacking H263 residue (Figure 1e, main text). Conversely, MGnify protein MGYP003574911459 (BLA09), was missed by PCM, deepARG, and CARD RGI strict, and was resistant to ampicillin when expressed in *E.coli* and contains the full active site architecture (Figure 1e, main text). Finally, a class D β-lactamase candidate k127_247171_3 and a dfr candidate k127_421717_1 from the lake metagenome (see Results) both functionally tested in this study share seqIDs of 59% and 41.5% to known ARGs, respectively. k127_247171_3 was only correctly predicted non-functional by ARG-PASS v2.0 and CARD RGI strict, whereas k127_421717_1 was incorrectly predicted functional by PCM and PLM-ARG. These examples suggest ARG-PASS v2.0 can more reliably distinguish functional from non-functional proteins across a range of seqIDs compared to existing methods.

The lowest sensitivity of ARG-PASS v2.0 was recorded for APH class, with 8 false negatives (FNs) identified (Table S3). After further interrogation, these 8 have closest homologs to APH(2”) enzymes. ARP structures encoded by APH(2”) form a cluster of 5 members at 20% seqID, thus with a sequence-structure distribution and decision boundary of reduced confidence and likely contributed to the non-functional prediction of these candidates. We anticipate that effects caused by clusters with a low number of members and thus training data will be reduced as ARG-PASS v2.0 is updated with ARGs identified in the future through complementary computational and functional metagenomic approaches.

#### CRS screen

We also analyzed the entire benchmarking dataset with the CRS search, to determine if these candidates would be identified by the CRS. Results for each candidate are presented as a column in Table S2. Altogether, we found the CRS had a recall of 81% for the functional candidates in the benchmarking dataset. Of note, no dfr or the AAC(2’) candidates were identified, which is likely due to the conserved regions of these sequences being too short, and continuous stretches of gaps which are too long to seed an alignment. As such, one solution here is to instead simply use the full-length of ARG sequences from these ARG classes for the alignment pre-screen with CRS and is the subject of ongoing work. In addition, a number of candidates from the PCM dataset were also missed (Class A β-lactamases, APH, class C β-lactamases). In this case, we cannot rule out the possibility that these genes had lower MICs due to testing differences by Ruppe et al. (2019), as discussed previously (Bartrop et al. 2026).

### Known ARGs identified in the doenjang metagenome

Among known candidates identified in the Doenjang metagenome by ARG-PASS v2.0, APH(6)-Id and APH(3″)-Ib (*strA* and *strB*) occurred adjacently as described by Gordon et al., (2008), and were detected in the UHGP-95 at ≥99% identity across globally distributed genomes including Asian isolates. *bla*_TEM-116_ was notably first reported from *K. pneumoniae* in Korean hospitals in 2004 (Jeong et al., 2004), and it was also identified in the doenjang metagenome, with a closest UHGP match of 97.9%, from *Vibrio cholerae* from Mozambique.

